# ZmCLA4 regulates leaf angle through multiple plant hormone-mediated signal pathways in maize

**DOI:** 10.1101/2020.03.20.999896

**Authors:** Dandan Dou, Shengbo Han, Lixia Ku, Huafeng Liu, Huihui Su, Zhenzhen Ren, Dongling Zhang, Haixia Zeng, Yahui Dong, Zhixie Liu, Fangfang Zhu, Qiannan Zhao, Jiarong Xie, Yajing Liu, Haiyang Cheng, Yanhui Chen

## Abstract

Leaf angle in cereals is an important agronomic trait contributing to plant architecture and grain yield by determining the plant compactness. Although ZmCLA4 was identified to shape plant architecture by affecting leaf angle, the detailed regulatory mechanism of ZmCLA4 in maize remains unclear. ZmCLA4 was identified as a transcriptional repressor using the Gal4-LexA/UAS system and transactivation analysis in yeast. The DNA affinity purification (DAP)-seq assay showed that ZmCLA4 not only acts as a repressor containing the EAR motif (CACCGGAC), but was also found to have two new motifs, CCGARGS and CDTCNTC. On analyzing the ZmCLA4-bound targeted genes, we found that ZmCLA4, as a cross node of multiple plant hormone-mediated pathways, directly bound to ARF22 and IAA26 to regulate auxin transport and mediated brassinosteroid signaling by directly binding to BZR3 and 14-3-3. ZmCLA4 bound two WRKY genes involved with abscisic acid, two genes (CYP75B1, CYP93D1) involved with jasmonic acid, B3 involved in the response to ethylene, and thereby negatively regulated leaf angle formation. We built a new regulatory network for the *ZmCLA4* gene controlling leaf angle in maize, which contributed to the understanding of ZmCLA4’s regulatory mechanism and will improve grain yields by facilitating optimization of plant architecture.

## Introduction

In the past few decades, improvement in plant architecture, especially the varieties with erect leaves, has played an important role in the genetic gain of maize yield (Lambert *et al*., 1978; Tian *et al.*, 2011). Upright leaves may have the additional benefit of reducing shade from neighboring plants, thereby increased planting density (Dubois and Brutnell, 2011). Therefore, it is an important selection target for ideal plant type breeding to have an upward leaf angle, and of great significance to clarify the genetic basis of the leaf angle for ideal maize breeding.

Research over the past several decades has identified quantitative trait loci (QTL) that regulate leaf angle by using bi-parental populations (Mickelson *et al.*, 2002; Lu *et al.*, 2007; Ku *et al.*, 2010, 2012; Chen *et al.*, 2015; Li *et al.*, 2015), and nested association mapping populations (Tian *et al.*, 2011) in maize. Taking advantage of 3D images from potted plants grown under greenhouse conditions, QTL analyses have been performed in maize using leaf angle data collected throughout the canopy at different developmental stages (McCormick *et al.*, 2016; Zhang *et al.*, 2017). QTLs explained between 0.45% and 85% of the phenotypic variance. Tian *et al.* (2019) mapped 12 QTLs for leaf angle in a population of 866 maize–teosinte BC_2_S_3_ recombinant inbred lines derived from a cross between the maize inbred line W22 and the teosinte accession CIMMYT 8759. To date, in the multiple QTLs that have been located, only six genes have been reported as a result of the combined use of quantitative genetics (Ku *et al.*, 2011; Zhang *et al*., 2014; Ren *et al.*, 2020; Tian *et al*., 2019; Cao *et al*., 2020). The first reported candidate gene, *ZmTAC1*, on a major QTL for LA that was detected on chromosome 2, was cloned using a comparative genomics method (Ku *et al.*, 2011). A nucleotide mutation in the 5′-untranslated region (UTR) influenced *ZmTAC1* expression (Ku *et al.*, 2011). UPA1 (Upright Plant Architecture1) and UPA2, two QTLs conferring upright plant architecture, were cloned by map-based cloning (Tian *et al*., 2019). UPA2 is controlled by a two-base sequence polymorphism that regulates the expression of a B3-domain transcription factor (ZmRAVL1) located 9.5 kilobases downstream. UPA2 exhibits differential binding by DRL1 (DROOPING LEAF1), and DRL1 physically interacts with *LG1* (LIGULELESS1) and represses LG1 activation of ZmRAVL1. ZmRAVL1 regulates brd1 (brassinosteroid C-6 oxidase1), which underlies UPA1, altering endogenous brassinosteroid content and leaf angle (Tian *et al.*, 2019). *ZmILI1*, a candidate gene of qLA2 on chromosome 2, directly binds to the promoters of the downstream gene *LG1* and activates *LG1* to further increase leaf angle (Ren *et al*., 2020). Ren *et al*. (2020) also found that *ZmILI1* and CYP90D1 formed a negative feedback loop to maintain the balance of brassinolide in maize. Cao *et al*. (2020) demonstrated that ZmIBH1-1 cloned by a map-based method negatively regulated leaf angle by causing cell wall lignification and cell elongation in the ligular region in maize.

In our previous study, ZmCLA4 cloned by a map-based method (Zhang *et al*., 2014) was identified as the ortholog of LAZY1 in rice and *Arabidopsis*. ZmCLA4 plays a negative role in the control of the maize leaf angle through the alteration of mRNA accumulation, leading to altered shoot gravitropism and cell development. In rice, LAZY1 regulates rice shoot gravitropism and tiller/leaf angle through an asymmetrical auxin pathway, which affects auxin transport, leading to its asymmetrical distribution (Li *et al*., 2007; Yoshihara and Iino, 2007). Zhang *et al*. (2018) demonstrated through transcriptome analysis that HSFA2D acts as a positive regulatory protein and plays a role upstream of LAZY1, and two transcription factors, WOX6 and WOX11, which are functionally redundant, play a role downstream of the LAZY1 signaling pathway in rice. Chen *et al.* (2011) showed that OsPIN2-overexpressing plants suppressed the expression of the gravitropism-related gene *OsLAZY1* in shoots, but did not alter the expression of OsPIN1b and OsTAC1, which were reported as tiller angle controllers. Li *et al*. (2019) showed that OsBRXL4 regulated shoot gravitropism by affecting polar auxin transport, similar to LAZY1. From the sequence analysis of AtLAZY1, Yoshihara *et al*. (2013) demonstrated that the functional domains inferred were nuclear localization signals, located between regions III and IV, and an EAR motif located in conserved region V. EAR motifs are often found in transcription regulators, and in many cases function as repressors (Kazan, 2006; Kagale *et al*., 2010). Although the identification and characterization of ZmCLA4 have increased our understanding of the role of gravitropism in shaping plant architecture, the detailed molecular mechanism involved in the regulation of polar auxin transport by LAZY1 remains unclear in plants. In this study, ZmCLA4 was identified as a transcriptional repressor using the Gal4-LexA/UAS system and transactivation analysis in yeast. The DNA affinity purification (DAP)-Seq assay showed that ZmCLA4, acting as a repressor, contained the EAR motif (CACCGGAC) and two new motifs (CCGARGS and CDTCNTC). Through the analysis of targeted genes bound by ZmCLA4, acting as a cross node of multiple plant hormone-mediated signaling pathways, ZmCLA4 not only directly bound to ARF22 and IAA26 to regulate auxin transport but also mediated brassinosteroid signal transduction by directly binding to BZR3 and 14-3-3. Additionally, it bound WRKYs involved with abscisic acid (ABA), two genes (CYP75B1, CYP93D1) involved with jasmonic acid (JA), and B3 involved in the response to ethylene, and thereby affected leaf angle formation. The results contributed to a better understanding of the ZmCLA4 regulatory pathways.

## Materials and methods

### Plant materials

The ZmCLA4-RNAi-transgenic (ZmCLA4-RT) line, with Yu368 as the genetic background, and the wild-type (WT) line (Zhang *et al.*, 2014) were sown in 2018. During the growth of the ZmCLA4-RT and WT lines, a sample from the leaf pulvinus in the 9-, 10-, and 11-leaf stages were obtained for each, immediately frozen in liquid nitrogen, and stored at −80°C before RNA/DNA extraction for molecular characterization and determination of the molecular mechanism of ZmCLA4.

### Transcriptional activation assay in yeast

The yeast strain YRG-2 (Stratagene, USA) containing the HIS3 and lacZ reporter genes was used to test transcriptional activation activity. The coding sequences (CDS) of ZmCLA4 were inserted into the pBD-GAL4 vector via EcoRI/SmaI sites, in which ZmCLA4 was fused with the GAL4 binding domain. The pBD-ZmCLA4, pBD-GAL4 (negative controls), and pGAL4 (positive control) plasmids were independently transfected into YRG-2 cells. The transfected yeast cells were grown on YPDA medium or SD/-Trp/-His medium for 3 d at 30 °C. β-Galactosidase filter assays were also conducted to determine the β-galactosidase activity of the transfected yeast cells by monitoring the generation of blue color, according to the method described in the Yeast Protocols Handbook (PT3024-1).

### Gal4/UAS System assay

35S-ZmCLA4 contains the cauliflower-mosaic virus (CaMV) 35S promoter, which drives ZmCLA4 expression. 35S-Luciferase contains firefly luciferase driven by the constitutive CaMV-35S promoter. The reporter gene constructs (UAS-GUS) and effector constructs (VP16, Gal4, and IAA17) were described previously by Tiwari *et al*. (2001). The ZmCLA4-GAL4 effector construct contains the full-length ZmCLA4 coding sequence fused to the N-terminus of the Gal4 DNA-binding domain under the control of the CaMV-35S promoter. The 35S-LUC construct was co-transformed as an internal control to normalize the GUS reporter gene expression. GUS and LUC enzymatic assays were performed in *Nicotiana benthamiana* leaves and performed according to Gampala *et al*. (2001).

### DAP-Seq experiments

DAP-Seq experiments were performed following the method described by O’Malley *et al.* (2016). First, a DAP-seq genomic DNA (gDNA) library was prepared by attaching a short DNA sequencing adaptor onto purified and fragmented gDNA. The adapter sequences were truncated Illumina TruSeq adapters; the TruSeq Universal and Index adapters corresponded to the DAP-seq Adapter A, CACGACGCTCTTCCGATCT, and Adapter B, GATCGGAAGAGCACACGTCTG. The DAP gDNA library was prepared using the kit from NEBNext® DNA Library Prep Master Mix Set for Illumina® (NEB #E6040S/L). ZmIBH1-1 was fused to HaloTag using the kit from pFN19K HaloTag T7 SP6 Flexi Vecto (cat#G184A) (Promega). ZmCLA4 fused to HaloTag was expressed using the TnT SP6 High-Yield Wheat Germ Protein Expression System (L3260) (Promega), and then purified using Magne HaloTag Beads (G7281) (Promega) and Magne HaloTag Beads. Next, the ZmCLA4-HaloTag mixture was incubated with 500 ng DNA library in 40 ul PBS buffer with slow rotation in a cold room for 1.5 h. The beads were washed five times with 200 μL PBS + NP40 (0.005%), and then were resuspended into the PBS buffer. The supernatant was removed and 25 μL of the EB buffer was added and incubated for 10 min at 98 °C to elute the bound DNA from the beads. The correct DAP-Seq library concentration to achieve a specific read count was calculated based on the library fragment size. Negative control mock DAP-Seq libraries were prepared as described above but without the addition of protein to the beads.

### DAP-Seq data analysis

We defined target genes as those that contained DAP-Seq peaks located within the transcribed regions of genes, in introns, 3 kb upstream of the transcription start site (TSS), or 3 kb downstream of the transcription termination site. DAP-Seq reads were aligned to the maize genome using Bowtie 2 (Langmead and Salzberg, 2012). Bowtie 2 supports the gapped and paired-end alignment modes. We ran Bowtie version 2.2.3 with default parameters and reported unique alignments. DAP-Seq peaks were detected using MACS2 (Zhang *et al*., 2008). We used MACS version 2.0.10, with default parameters because duplicates were allowed, with the q-value < 0.05.

### Electrophoretic mobility shift assay

The full-length ZmCLA4 cDNA was amplified with gene primers (Supplementary Table S1) and fused into the *Sgf*I and *Pme*I sites of pFN19K HaloTag^®^ T7 SP6 Flexi^®^ Vector. The HaloTag-CLA4 fusion protein was expressed using the TNT^®^ Coupled Wheat Germ Extract Systems (Promega, Fitchburg, WI, USA) and Magne^®^ HaloTag Beads (Promega) EMSA. Oligonucleotide probes (Supplementary Table S1) were synthesized and labeled according to the standard protocol of Invitrogen Technology (Shanghai, China). We used standard reaction mixtures for the EMSA containing 20 ng of purified ZmCLA4 fusion protein, 5 ng of biotin-labeled annealed oligonucleotides, 2 μL of 10× binding buffer (100 mM Tris, 500 mM KCl, and 10 mM DTT, pH 7.5), 1 μL of 50% (v/v) glycerol, 1 μL of 100 mM MgCl2, 1 μL of 1 mg mL^−1^ poly(dI– dC), 1 μL of 1% (v/v) Nonidet P-40, and double-distilled water to obtain a final volume of 20 μL. The reactions were incubated at 25 °C for 20 min, electrophoresed in 6% (w/v) polyacrylamide gels, and then transferred to N+ nylon membranes (Millipore, Darmstadt, Germany) in 0.53× TBE (Tris-Borate–EDTA) buffer at 380 mA and 4 °C for 30 min. Biotin-labeled DNA was detected using the LightShift™ Chemiluminescent EMSA kit (Thermo Fisher). Bands were visualized using the Chemiluminescent Western Blot Detection Kit (Thermo Fisher).

### Transient assays for *in vivo* activation activity

To generate the Pro CLA4, luciferase (LUC) reporters for the dual-luciferase assays, ∼2500 bp from the TSS promoter region of potential targets for stress response, were inserted into pGreenII0800-LUC. To generate the CaMV 35S promoter-driven ZmCLA4 effector, the full-length coding sequence of ZmCLA4 was inserted into pUC18-35S. Transient dual-luciferase assays were performed in *N. benthamiana* leaves and checked using dual-luciferase assay reagents (Promega). Following infiltration, plants were maintained at room temperature under a 14/10 h light/dark photoperiod. Leaf protein was extracted 48 h later. The protein was extracted using a passive lysis buffer (Cat# E1910, Promega). LUC activity was measured using a GloMax®20/20 Luminometer (Cat# E5311, Promega). Then, 100 µL of Stop and Glow Buffer was added to the reaction and Renilla luciferase (REN) activity was measured. For this analysis, the ratio between LUC and REN activities was measured three times.

### Real-time reverse transcription-PCR (qRT-PCR)

Total RNA was isolated from the collected samples using TRIzol reagent (Invitrogen, Waltham, MA, USA) and treated with RNase-free DNase I to remove DNA contamination. cDNA was synthesized using an M-MLV reverse transcriptase-based cDNA first-strand synthesis kit (Invitrogen). qRT-PCR was performed using the SYBR^®^ Green CR Master Mix Kit (Applied Biosystems, Waltham, MA, USA), following the manufacturer’s protocol on a LightCycler^®^ 480II Sequence Detection System. Relative gene expression was calculated according to the 2^−ΔΔCt^ method. Expression values were normalized to the *18S* ribosomal gene for qRT-PCR. The primer sequences used in the qRT-PCR assay are listed in Supplementary Table S1.

## Results

### ZmCLA4 acts as a transcription repressor

A previous study identified ZmCLA4 with some sequence similarity to the rice gene OsLAZY1 and Arabidopsis gene AtLAZY1 (Li *et al*., 2007; Dong *et al*., 2013; Yoshihara *et al*., 2013; Zhang *et al*., 2014). The only functional domain inferred from the sequence analysis of ZmCLA4 was an ethylene-responsive element binding factor-associated amphiphilic repression (EAR) motif located in the conserved region (Figure 1A) and was frequently identified at the C-terminal end of the protein (Ohta *et al*., 2001). EAR motifs are often found in transcription regulators, and in many cases, function as repressors (Kazan, 2006; Kagale *et al*., 2010). To confirm that ZmCLA4 can activate or inhibit gene expression as a transcription factor, ZmCLA4 was first subjected to transactivation analysis in yeast. The results showed that ZmCLA4 had no transcriptional activation activity in yeast (Figure 1B). We then used the Gal4-LexA/UAS system, which tests ZmCLA4 for positive or negative transcriptional potential (Tiwari *et al*., 2001). The reporter gene was expressed at high levels when co-transformed into protoplasts with the LexAVP16 fusion construct, which contained the coding sequences for the LexA DNA-binding domain fused in frame with the coding sequence of the VP16 transcriptional activation domain. As reported previously (Tiwari *et al*., 2001), Gal4 fusion with the transcriptional repressor domain of IAA17 (IAA17a1) strongly reduced the expression of the reporter gene. Similarly, the co-transformation of the reporter gene with a construct for the ZmCLA4-Gal4 fusion protein (ZmCLA4-Gal4) significantly decreased reporter gene expression (Figure 1C). Together, these results demonstrate that ZmCLA4 is a transcriptional repressor.

**Fig. 1.**
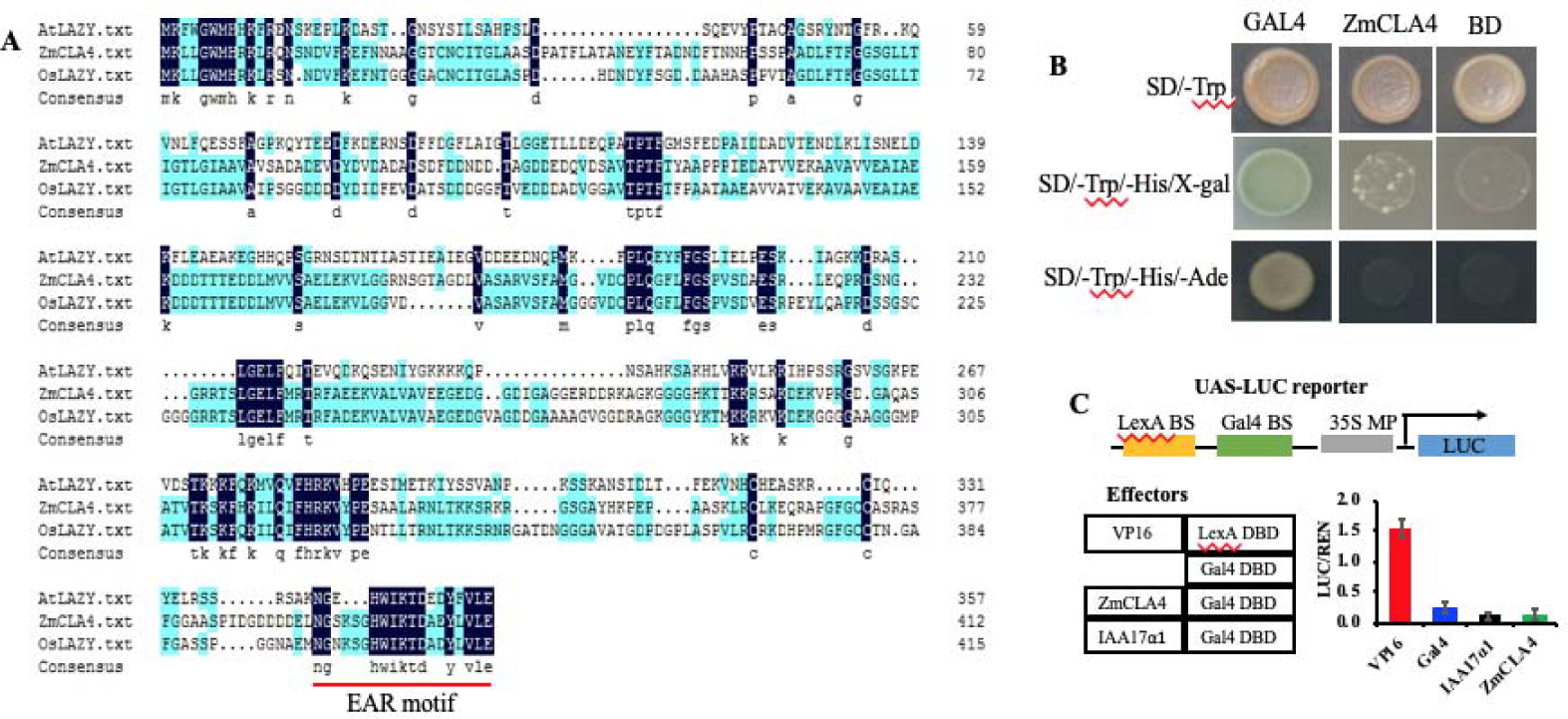
Characteristic analysis of ZmCLA4 protein. **(A)** Sequence alignment of maize CLA4, rice LAZY1, and Arabidopsis AtLAZY1 amino acid sequences. The alignment was performed using DNAMAN. Identical amino acids are in black and the conserved EAR motif is underlined in red. **(B)** Transactivation analysis of ZmCLA4 fused to the GAL4 DNA-binding domain in yeast. **(C)** Transient assays of the transcriptional activity of ZmIBH1-1. *Nicotiana benthamiana* leaves were transformed with the reporter (UAS-LUC) and effector constructs (left), and the reporter gene expression was determined (right). Data are means (± SD), n = 3. UAS-LUC, reporter construct containing Gal4 and LexA binding sites, and a 35S minimal promoter upstream of the coding sequence of LUC; VP16, VP16 fused to the LexA DNA-binding domain (DBD); Gal4, Gal4 DBD; IAA17a1, the transcription repression domain of IAA17 fused to the Gal4 DBD; ZmCLA4, full-length ZmCLA4 fused to Gal4 DBD. The LUC reporter gene expression was normalized to the luciferase activity and presented as values relative to the VP16 control, the value of which was set as 1.

### DAP-Seq identifies genes directly targeted by ZmCLA4

To investigate the regulatory mechanism mediated by *ZmCLA4*, we performed a DAP-Seq assay to uncover the genes directly targeted by ZmCLA4. Using the Illumina platform (50 bp-long pair-end reads), the DAP-Seq assay produced ∼2.8 million reads for each sample, of which ∼2.1 million reads were uniquely mapped to the maize_V4 genome sequence with an effective read ratio of ∼75% (Table S2). We predicted ZmCLA4 binding sites using the software MACS2 (Zhang *et al*., 2008) with a p-value < 0.05 (based on a Poisson distribution comparing the ZmCLA4 sample and the control) and identified 7182 peaks across the entire genome through the ZmCLA4 binding motifs CACCGGAC, CCGARGS, and CDTCNTC (Figure 2A). Of the ZmCLA4 binding sites, 49.44% (3524 peaks) were located in the genic regions containing the genes, as well as 3 kb upstream of the start codon and 3 kb downstream of the stop codon (Figure 2B, C). Among these genic region peaks, 35.62%, 1.25%, 12.34%, 14.63%, and 36.16% are located in the promoter, 5′ -UTRs, introns, exons, and the transcription termination region, respectively.

**Fig. 2.**
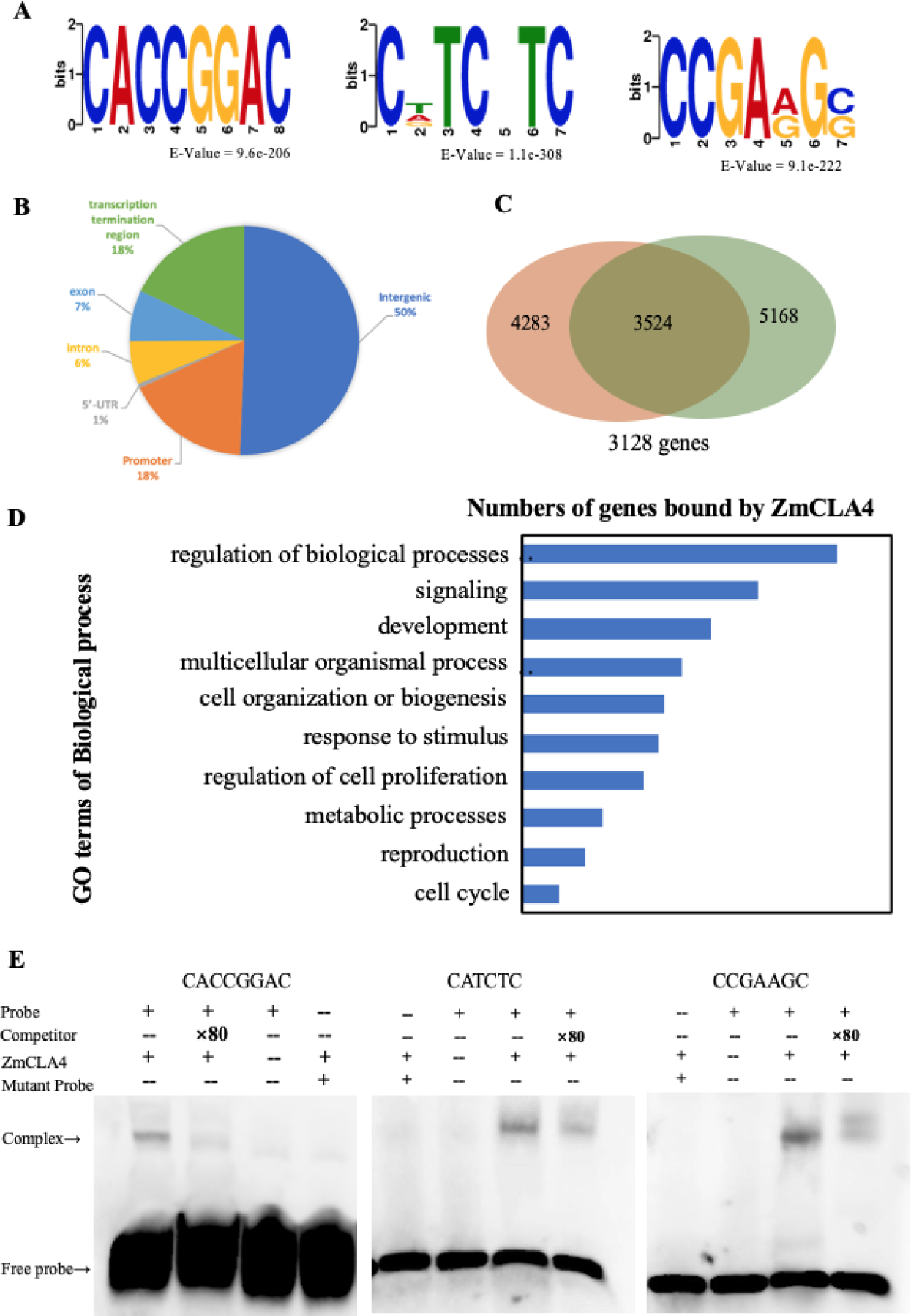
DAP-seq analysis of maize *ZmCLA4.* **(A)** ZmCLA4 binding to CACCGGAC, CCGARGS, and CDTCNTC motifs as identified by the MEME-ChIP. **(B)** Distribution of the ZmCLA4 binding sites. **(C)** Comparison of ZmCLA4-occupied peaks using two biological replicates. **(D)** GO annotation of targeted genes bound by ZmCLA4. The y-axis represents the percentage of genes related to each functional category. **(E)** Results of EMSAs confirming ZmCLA4 binding to CACCGGAC, CCGARGS, and CDTCNTC.

The 3524 peaks corresponded to 3128 genes. Stringent Gene Ontology (GO) term enrichment analysis of the 3128 genes revealed that ZmCLA4 binding genes are mainly involved in the regulation of biological processes, multicellular organismal processes, development, signaling, responses to stimuli, and regulation of cell proliferation, among others (Figure 2D). More specifically, ZmCLA4 binds to the upstream region of 1114 genes involved in biological regulation, signaling, regulation of cell proliferation, and response to stimuli. Of the 1114 genes bound to the upstream regions, 16 genes appeared to be responsible for the leaf angle. These genes were involved in response to phytohormones, such as ABA, auxin, and brassinolide (Table S3).

### Binding Motif Analysis Revealed Novel ZmCLA4 cis-Elements

To confirm that ZmCLA4 was bound to the predicted EAR cis-element (CACCGGAC), an EMSA was performed using a purified ZmCLA4 protein and a labeled DNA probe containing the ZmCLA4-binding site (CACCGGAC). As shown in Figure 2E, ZmCLA4 bound to CACCGGAC. The addition of an 80× unlabeled competitor reduced the detected binding of ZmCLA4, and it did not bind to mutant probes (CACCGGAC mutated into TCTTTTGT). Without the ZmCLA4 protein, no bands were observed, except for the free probe. These results further confirmed the specific binding of ZmCLA4 to CACCGGAC.

To explore novel ZmCLA4 binding motifs, the 3 kb flanking sequences around all of the genic peak summits were applied to the motif discovery tool MEME-ChIP. The motifs CCGARGS and CDTCNTC were identified as statistically defined motifs (E-Value = 9.1 e-222, E-Value = 1.1 e-308, respectively; Figure 2A). Cis-element scanning was performed using the CCGARGS and CDTCNTC motifs on the 3 kb flanking sequences around the peak maxima, and 1086 potential ZmCLA4 directly targeted genes with functional annotations were detected. Among the 1086 genes, 195 were in the promoter regions and combined with CCGARGS and CDTCNTC. Confirming the labeled DNA probe contained the ZmCLA4-binding sites, CCGARGS and CDTCNTC, Figure 2E shows ZmCLA4 bound to CCGAAGC and CTTCGTC. Furthermore, the addition of the 80× unlabeled competitor reduced the detected binding of ZmCLA4, and it did not bind to mutant probes (CCGAAGC and CTTCGTC mutated into TTCTTAT and AACGTTAGT, respectively). Without the ZmCLA4 protein, no bands were observed, except for the free probe. These results confirmed the specific binding of ZmCLA4 to CCGAAGC and CTTCGTC.

### ZmCLA4 regulates auxin transport by directly binding to ARF and IAA transcription factors in maize

Li *et al.* (2007) and Yoshihara *et al.* (2007) demonstrated that OsLAZY affects rice shoot gravitropism and tiller/leaf angle by negatively regulating auxin transport, leading to the asymmetric distribution of auxin. ZmCLA4, an ortholog of OsLAZY, was identified to directly bind to auxin response factors (Zm00001d036593, ARF22; Zm00001d039513, IAA26; Table S3). To understand whether ZmCLA4 functions as a transcriptional repressor of the auxin-respective genes, we performed a dual-luciferase transient transcriptional activity assay using *N. benthamiana* leaves with ZmCLA4 driven by the cauliflower-mosaic virus (CaMV) 35S promoter as an effector and LUC (the firefly luciferase-coding gene driven by −500-to −3000 auxin-respective genes) as the reporter gene (Figure 3A). The results showed that ZmCLA4 specifically repressed the expression of LUC with the ARF22 promoter and increased the expression of LUC with the IAA26 promoter, indicating that the two genes are target genes of ZmCLA4 (Figure 3B). Additionally, we found that ZmCLA4 binds to three ABC transporter G family members (Zm00001d042953, ABC-2; Zm00001d009243, MRP10; Zm00001d023392, ABCG37; Table S3) mediating the efflux of the auxin precursor. The results showed that ZmCLA4 specifically repressed the expression of LUC with the ABC-2 promoter, MRP10 promoter, and ABCG37 promoter, indicating that these genes are the target genes of ZmCLA4 (Figure 3B).

**Fig. 3.**
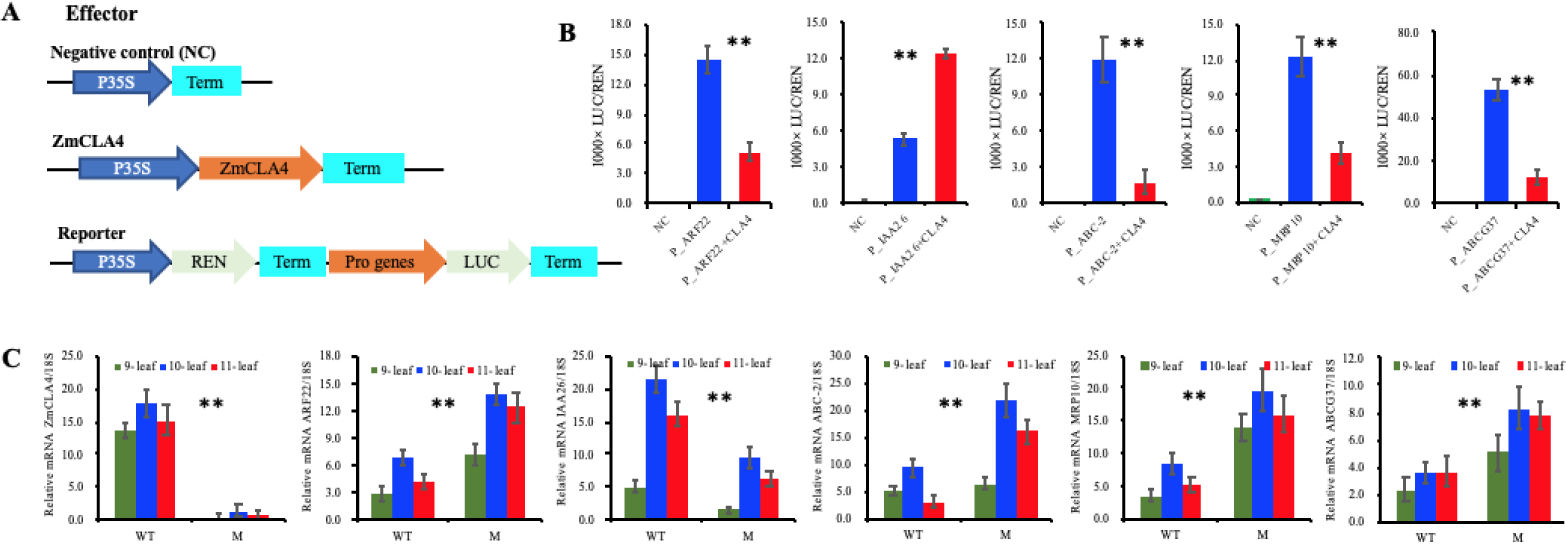
ZmCLA4 functions as a transcriptional repressor of the genes involved in response to auxins. **(A)** The 35S:REN-Pro PR:LUC reporter constructs were transiently expressed in *Nicotiana benthamiana* leaves together with the control vector or 35S: ZmCLA4 effector. **(B)** The LUC/REN ratio represents the relative activity of the gene promoters (*P < 0.05, **P < 0.01). **(C)** Expression analysis of the targeted genes in wild-type and ZmCLA4-RNAi plants using leaf pulvini from the 9-leaf, 10-leaf, and 11-leaf stages.

To understand the effects of ZmCLA4 on ARF22, IAA26, ABC-2, MRP10, and ABCG37 *in vivo*, we measured mRNA levels of the five genes by qRT-PCR in WT and ZmCLA4-RNAi plants. The ZmCLA4 was expressed in the WT but not in ZmCLA4-RNAi plants in leaf pulvinus of 9-leaf, 10-leaf, and 11-leaf, whereas mRNA levels of ARF22, ABC-2, MRP10, and ABCG37 were significantly higher, and the mRNA level of IAA26 was significantly lower in the leaf pulvini of the ZmCLA4-RNAi plants than in WT plants (Figure 3B). These results were consistent with the expression results of LUC with corresponding gene promoters.

### ZmCLA4 mediates brassinosteroid signal transduction by directly binding to BZR3 and 14-3-3 in maize

Among the genes bound by ZmCLA4, there are two genes (Zm00001d053543 and BZR3; Zm00001d003401, 14-3-3; Table S3) involved in brassinosteroid signal transduction. We performed dual-luciferase transient transcriptional activity assays with the ZmCLA4 protein and LUC driven by the promoter sequences of the two genes as reporters (Figure 3A). The results showed that ZmCLA4 specifically repressed the expression of LUC with the BZR3 promoter and increased the expression of the 14-3-3 promoter, indicating that the two genes are target genes of ZmCLA4 (Figure 4A).

**Fig. 4.**
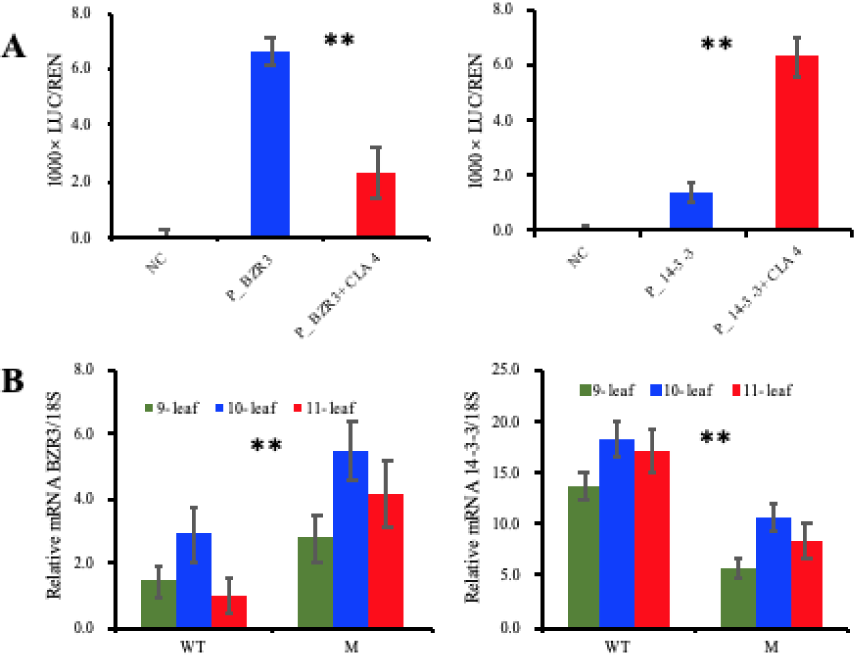
ZmCLA4 functions as a transcriptional repressor of the genes involved in response to brassinolide. **(A)** The LUC/REN ratio represents the relative activity of the gene promoters (*P < 0.05, **P < 0.01). **(B)** Expression analysis of the targeted genes involved in responses to brassinolide in wild-type and ZmCLA4-RNAi plants using leaf pulvini from the 9-leaf, 10-leaf, and 11-leaf stages.

To further understand the effects of ZmCLA4 on BZR3 and 14-3-3 *in vivo*, we measured mRNA levels of the two genes by qRT-PCR in WT and ZmCLA4-RNAi plants. The mRNA level of BZR3 was significantly higher in leaf pulvini of the ZmCLA4-RNAi plants than in the WT in the 9-leaf, 10-leaf, and 11-leaf. The mRNA level of 14-3-3 was significantly lower in leaf pulvini of the ZmCLA4-RNAi plants than in WT plants in the 9-leaf, 10-leaf, and 11-leaf (Figure 4A). These results were consistent with the expression results of LUC with corresponding gene promoters.

### ZmCLA4 directly regulates other phytohormone-respective genes

In this study, we also found that ZmCLA4 binds to four genes (Zm00001d035167, GTE8; Zm00001d050723, SnRK2.6; Zm00001d013240, WRKY4; Zm00001d003331, WRKY72) in response to ABA, two genes (Zm00001d047425, CYP75B1; Zm00001d046758, CYP93D1) associated with JAs, two genes (Zm00001d024545, B3; Zm00001d007522, ETO1) involved in the response to ethylene. We performed dual-luciferase transient transcriptional activity assays to investigate whether ZmCLA4 could function as a repressor of these genes. The results showed that LUC expression driven by GTE8, SnRK2.6, WRKY4, WRKY72, CYP75B1, CYP93D, B3, and ETO1 promoters was prominently induced by ZmCLA4 (Figure 5A), indicating that these genes are the target genes of ZmCLA4.

**Fig. 5.**
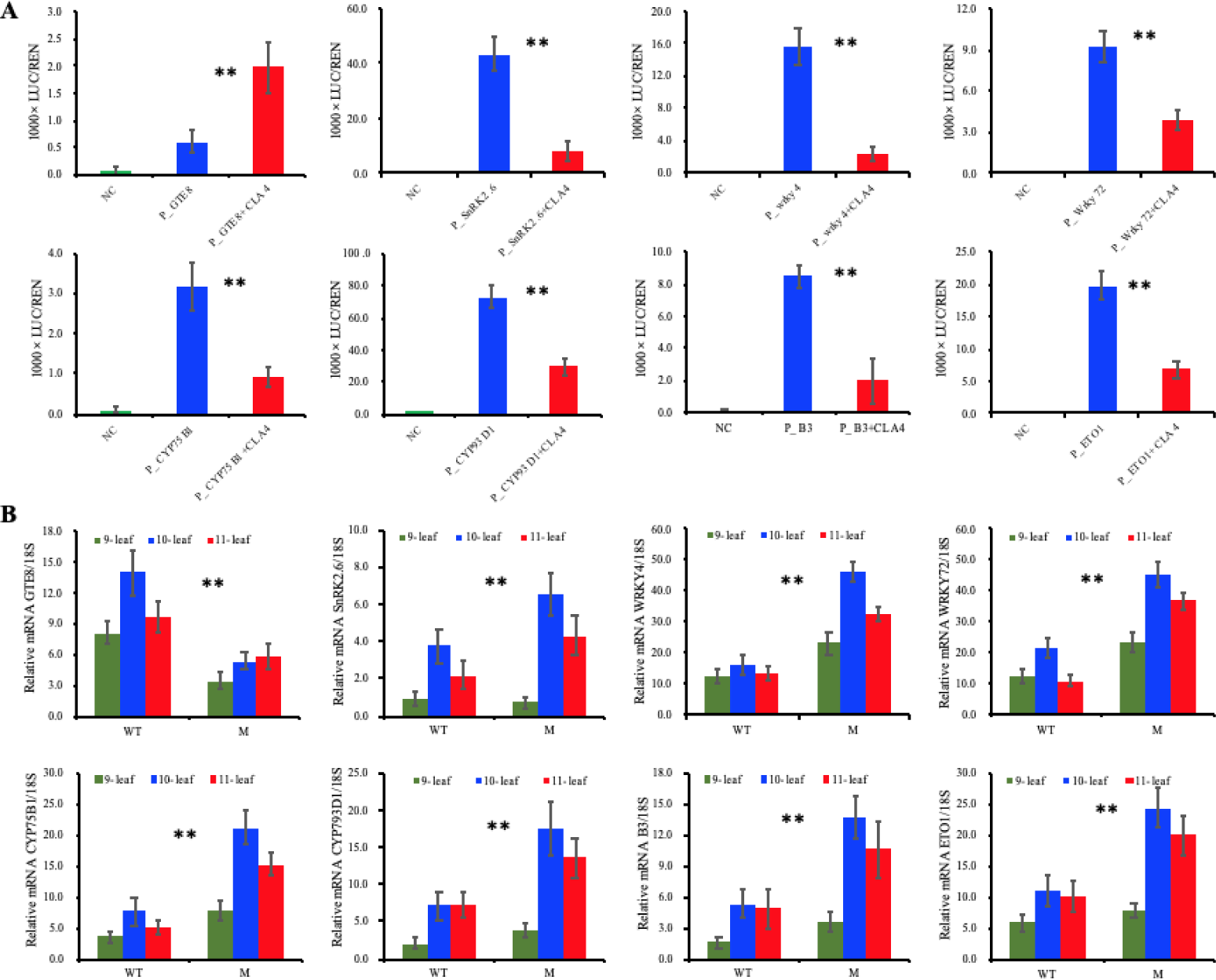
ZmCLA4 functions as a transcriptional repressor of other phytohormone-related genes. **(A)** The LUC/REN ratio represents the relative activity of the gene promoters (*P < 0.05, **P < 0.01). **(B)** Expression analysis of the other phytohormone-respective targeted genes in wild-type and ZmCLA4-RNAi plants using leaf pulvini from the 9-leaf, 10-leaf, and 11-leaf stages.

**Fig. 6.**
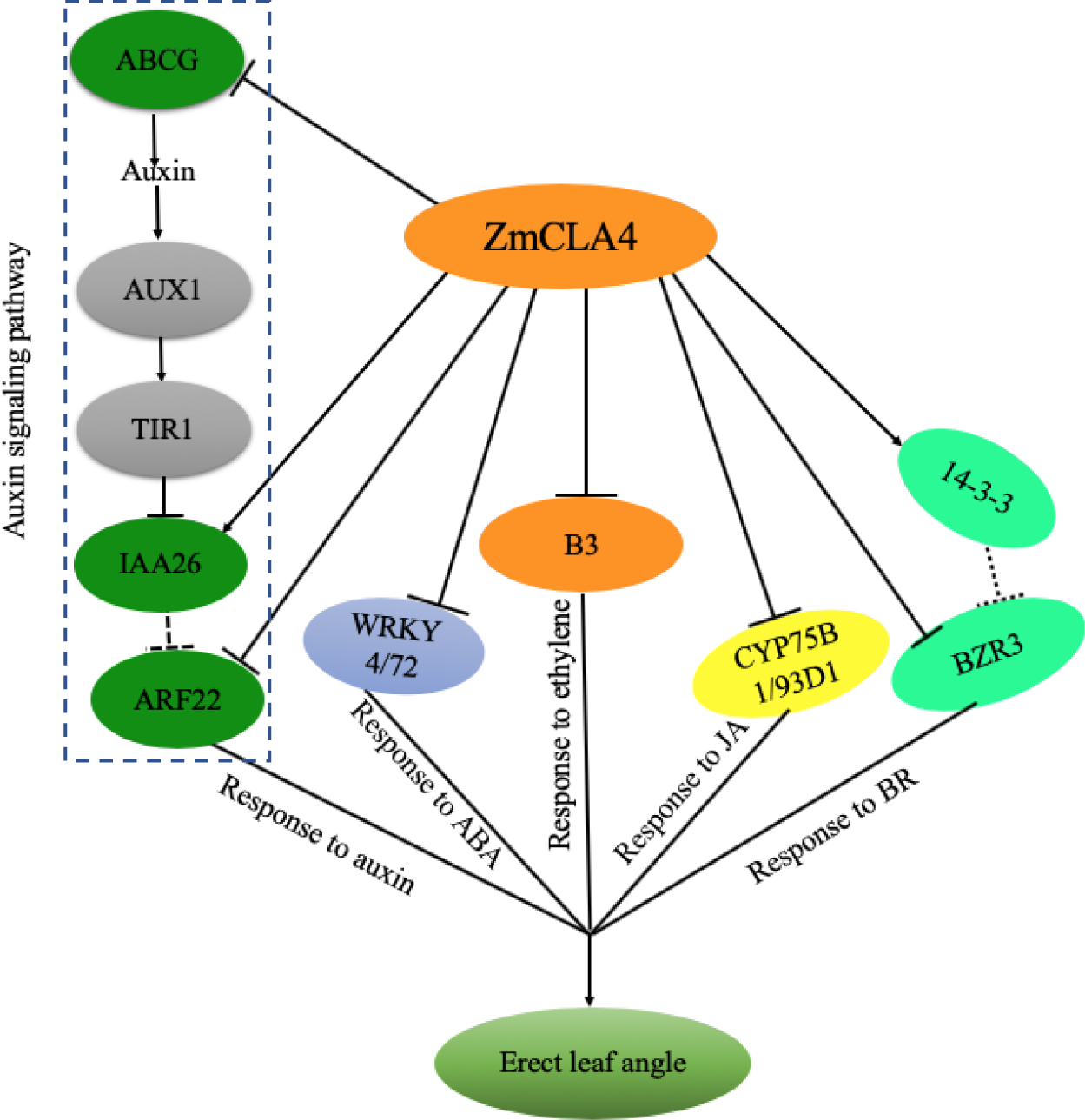
A schematic model for leaf angle formation in maize. The arrows between the genes represent promotion or activation, and the ┴ bars between the genes indicate suppression. The green circles represent auxin-responsive genes, the yellow circle represents the jasmonic acid-responsive gene, the bright green circles represent the brassinosteroid-responsive genes, the orange circles represent ethylene-responsive genes, and the lavender circle represents the abscisic acid-responsive genes.

To further understand the effects of ZmCLA4 on GTE8, SnRK2.6, WRKY4, WRKY72, CYP75B1, CYP93D, B3, and ETO1 *in vivo*, we measured the mRNA levels of the eight genes by qRT-PCR in WT and ZmCLA4-RNAi plants. The mRNA levels of SnRK2.6, WRKY4, WRKY72, CYP75B1, CYP93D, B3, and ETO1 were significantly higher in leaf pulvini of the ZmCLA4-RNAi plants than in the WT in the 9-leaf, 10-leaf, and 11-leaf, and the mRNA level of GTE8 was significantly lower in leaf pulvini of the ZmCLA4-RNAi plants than in the WT in the 9-leaf, 10-leaf, and 11-leaf (Figure 5B). These results were consistent with the expression results of LUC with corresponding gene promoters.

## Discussion

Leaf angle is a key agronomic trait determining maize plant architecture and grain yield per unit area. Maize plant architecture with more upright leaves (i.e., smaller leaf angle) decreases mutual shading and sustains light capture for photosynthesis; thus, increasing the accumulation of leaf nitrogen for grain filling and increasing grain yield. ZmCLA4 plays a negative role in the control of maize LA through the alteration of mRNA accumulation. However, the regulatory mechanism of ZmCLA4 has not been reported in maize. In this study, sequence analysis showed that ZmCLA4 acts as a repressor including an EAR motif located in the conserved region. First, we identified that ZmCLA4 had no transcriptional activation activity through transactivation analysis in yeast. Then, ZmCLA4 was identified as a transcriptional repressor through the Gal4-LexA/UAS system. The EAR motif was located at the C-terminal end of the AtLAZY1 protein (Yoshihara *et al*., 2013), whereas the conclusion was not proved by corresponding experiments in *Arabidopsis*. The results of the DAP-Seq assay showed that ZmCLA4 directly affected the expression of target genes through binding motifs CACCGGAC (EAR motif), CCGARGS, and CDTCNTC. This result not only further proved that ZmCLA4 acts as a repressor containing the EAR motif, but also identified two new motifs (CCGARGS and CDTCNTC), thereby providing a solid basis for further analysis of the molecular characteristics of ZmCLA4.

In rice, OsLAZY1 plays a negative role in polar auxin transport (PAT), and loss-of-function of OsLAZY1 greatly enhances PAT; thus, altering the endogenous IAA distribution in shoots, leading to reduced gravitropism, and the leaf/tiller-spreading phenotype of rice plants (Li *et al*., 2007). However, the detailed molecular mechanism involved in the regulation of PAT by LAZY1 is still unclear. In this study, ZmCLA4, an ortholog of OsLAZY, directly bound to auxin response factors ARF22 and IAA26 to participate in the auxin signal transduction pathway TIRI/AFB-Aux/IAA/TPL-ARFs, thereby affecting PAT. The auxin receptor TIR1 and its homologous receptor protein AFBs interact with Aux/IAA protein, which is an inhibitor of auxin signaling. Then, auxin stabilizes TIRI/AFB and Aux/IAA protein interaction, and degrades Aux/IAA protein, thus releasing ARFs, which are inhibited by the Aux/IAA protein, to mediate auxin signal transduction (Dharmasiri *et al*., 2005 *a*, b; Kepinski and Leyser, 2005 *a, b*). ARFs are important for the normal growth and development of plants. In *Arabidopsis*, they control leaf development. ARF5 is critical to leaf initiation and vein pattern formation (Garrett *et al*., 2012). OsARF19-overexpression lines show an enlarged lamina inclination (leaf angle increase) because of an increase in its adaxial cell division in rice (Zhang *et al*., 2015). Our qRT-PCR results showed that the mRNA level of ARF22 was significantly higher and that of IAA26 was significantly lower in the leaf pulvini of the ZmCLA4-RNAi plants than in WT plants. Additionally, ZmCLA4 directly bound to three ABC transporter G family members (ABC-2, MRP10, ABCG37), mediating the efflux of the auxin precursor. The study showed that ABCG36 mediates the efflux of the auxin precursor indole 3-butyric acid (IBA) from roots, as evidenced by hypersensitive root growth phenotypes of *abcg36* mutants in the presence of IBA or precursors of synthetic auxin analogs in Arabidopsis (Strader *et al*., 2009; Ru□z□ic□ka *et al*., 2010). The function as a transporter of IBA and various auxinic compounds has also been assigned to ABCG37/PDR9, as evidenced by altered responsiveness of *abcg37* mutant plants to synthetic auxins and inhibitors of auxin transport, but not to IAA, the endogenous auxin (Ru□z□ic□ka *et al*., 2010; Ito and Gray, 2006; Strader *et al*., 2008). In this study, the qRT-PCR results showed that mRNA levels of ABC-2, MRP10, and ABCG37 were significantly higher in leaf pulvini of the ZmCLA4-RNAi plants than in WT plants. These results implied that ZmCLA4 could negatively regulate PAT by directly repressing ABC transporter G family members. Overall, these results suggest that ZmCLA4 represses ARF22 by directly promoting IAA26 or directly repressing ABCGs or ARF22 and then mediated auxin signal transduction, and finally led to a decrease in leaf angle.

The signaling pathways of various hormones in plants often cross each other, forming a complex regulatory network. In terms of auxin, the interaction between BR, ABA, JA, and other hormones has been extensively studied. In rice, OsLAZY1 plays a negative role in PAT (Li *et al*., 2007). Except for ZmCLA4 directly binding to ARF22 and IAA26 to participate in the pathway, ZmCLA4 not only mediates brassinosteroid signal transduction through directly binding to BZR3 and 14-3-3 in maize, but also binds to four genes (GTE8, SnRK2.6, WRKY4, WRKY72) in response to ABA, two genes (CYP75B1 and CYP93D1) associated with JA, and two genes (B3, ETO1) involved in the response to ethylene. In rice, the BR transcription factor BZR1 increases leaf angle, whereas RNAi:BZR1 plants have leaves with small inclinations (Bai *et al*., 2007; Zhang *et al*., 2012). BZR3, as a homologous gene of BZR1, could increase the leaf angle in maize. BRs are a group of steroid hormones with a paracrine mode of action that determines important traits, such as plant architecture and have been extensively reported as key regulators of leaf angle in cereals. This conclusion is derived from the numerous BR biosynthesis or signaling mutants investigated in rice, maize, and sorghum with consistently reduced leaf angles (Yamamuro *et al.*, 2000; Morinaka *et al.*, 2006; Sakamoto *et al.*, 2006; Wang *et al.*, 2008; Divi and Krishna, 2009; Makarevitch *et al.*, 2012; Tong *et al.*, 2014; Sun *et al.*, 2015; Best *et al.*, 2016; Feng *et al.*, 2016; Hirano *et al.*, 2017). The study of the BR signaling pathway in Arabidopsis and rice is clear; that is, BR is sensed by BRI1 and its two homologous proteins BRL1 and BRL3. The unphosphorylated BKI1 interacts with BRI1, prevents the interaction between BRI1 and BAK1, and inhibits the activation of BR signaling (Jiang *et al.*, 2015). BKI1 also competitively binds to the 14-3-3 protein that assists in the degradation of BZR1, reducing the negative regulatory effect of the 14-3-3 protein on BZR1, and then rapidly promotes the transmission of the BR signal (Wang *et al*., 2011). Based on our data, mRNA levels of BZR3 were significantly higher in ZmCLA4-RNAi plants than in the WT, and mRNA levels of 14-3-3 were significantly lower in ZmCLA4-RNAi plants than in WT plants, indicating that ZmCLA4 was involved in BRI1/BAK1 mediated BR signaling by directly binding BZR3 and 14-3-3 to regulate leaf angle in maize.

Early evidence suggests that ABA reduces leaf angle and inhibits the action of externally applied BRs (Wada *et al*., 1981). WRKYs include ABA-responsive elements and might function as positive regulators in mediating plant responses to ABA (Jiang and Deyholos, 2009; Gao *et al*., 2011). Accumulating evidence has revealed that WRKY proteins play diverse roles in responses to biotic and abiotic stresses and are involved in various processes of plant growth and development by regulating the expression of target genes via binding to the W-box cis-element (Rushton *et al*., 2010). For example, OsWRKY53 overexpression led to enlarged leaf angles and increased grain size, in contrast to the erect leaves and smaller seeds in the oswrky53 mutant (Tian *et al*., 2017). Based on our data, the mRNA levels of WRKY4 and WRKY72 were significantly higher in ZmCLA4-RNAi plants than in WT plants, indicating that ZmCLA4 directly repressed the expression of ABA-responsive WRKY4/72 (homologous gene of OsWRKY53) and mediated the ABA pathway to reduce leaf angle in maize.

JAs also regulate leaf angles through their interaction with BR metabolism. Methyl-JA represses the expression of BR biosynthesis and signaling genes, reducing endogenous levels of BRs (Gan *et al.*, 2015), and thereby, leaf angle. CYP75B1 and CYP93D1 were both annotated as “response to jasmonic acid (GO:0009753).” Heitz *et al*. (2012) showed that cytochrome P450, CYP94C1, and CYP94B3 are involved in JA-Ile oxidation. CYP93G1 converts naringenin to apigenin, and then CYP75B4 converts apigenin to luteolin, which is further metabolized to tricin through *O*-methyltransferase activity and the chrysoeriol 5′-hydroxylase activity of CYP75B4 in rice (Lam *et al*., 2015; Sangkyu *et al*., 2016). Tricin derivatives have been reported to be incorporated into lignin (Li *et al*., 2016). These results imply that ZmCLA4 directly regulates CYP75B1 and CYP93D1 (homologous genes of CYP75B and CYP93G, respectively) to mediate the JA pathway and affect lignin biosynthesis, thereby affecting the leaf angle in maize.

Ethylene is a gaseous plant hormone that plays a key role in leaf senescence (Abeles *et al*., 1992). BR-induced rice lamina joint inclination was accompanied by increased ethylene production because of greater expansion of the adaxial cells relative to the dorsal cells in the lamina joint (Cao and Chen, 1995). We also found that ZmCLA4 binds two genes (B3 and ETO1) involved in response to ethylene. In particular, B3 encodes a B3 domain containing a protein homologous to ZmRAVL1 in maize (Tian *et al*., 2019). ZmRAVL1 RNAi and knockout lines exhibited smaller leaf angles in the lower, middle, and upper leaves compared with that of the WT plants by increasing adaxial sclerenchyma cells in the ligular region (Tian *et al*., 2019). From our data, B3 was significantly higher in ZmCLA4-RNAi plants than in WT plants. These results showed that ZmCLA4 decreased the leaf angle by directly repressing the expression of ethylene-responsive B3 in maize.

In summary, our results demonstrated that ZmCLA4, a cross node of five major phytohormone signaling pathways, negatively regulates leaf angle in maize. It directly affects the patterns of gene transcription. We built a hierarchical regulatory model describing ZmCLA4 and ZmCLA4-targeted genes to explain the effects on the formation of leaf angle during maize development. Our data suggest that ZmCLA4 directly represses genes associated with a range of biological responses, including auxin, BR, ABA, JA, and ethylene signaling pathways, resulting in an erect leaf angle. The manipulation of the regulation of leaf angle in maize reported herein, to adapt to high-density planting, has provided important insights that will help in directing future approaches for the production of high-yield maize varieties. A better understanding of ZmCLA4 regulatory pathways will provide new insights into the effects of ZmCLA4 through multiple signaling pathways in the future.

## Acknowledgements

This research was supported by grants from the Central Plains Science and Technology Innovation Leading Talents (194200510021), National Key Research and Development Program of China (2016YFD0101803) and National Natural Science Foundation of China (No. 31871639).

## Author contributions

D.D., S. H. Z. R., D. Z., H. Z., Y. D., Z. L., F. Z., Q. Z., J. X. H. L. and H. C conducted molecular biology experiments; D. D. and H. S. analyze the data. Y. C, L.K., D. D. and S. H. designed the experiments and wrote the manuscript.

## Supporting information

The following materials are available in the online version of this article.

**Table S1. Primer sequences used for the experiments.**

**Table S2. The summary of reads analysis.**

**Table S3. CLA4 regulating leaf angle-related genes in maize.**

